# Somatic Truth Data from Cell Lineage

**DOI:** 10.1101/825042

**Authors:** Megan Shand, Jose Soto, Lee Lichtenstein, David Benjamin, Yossi Farjoun, Yehuda Brody, Yosef E. Maruvka, Paul C. Blainey, Eric Banks

## Abstract

Existing somatic benchmark datasets for human sequencing data use germline variants, synthetic methods, or expensive validations, none of which are satisfactory for providing a large collection of true somatic variation across a whole genome. Here we propose a dataset of short somatic mutations, that are validated using a known cell lineage. The dataset contains 56,974 (2,687 unique) Single Nucleotide Variations (SNV), 6,370 (316 unique) small Insertions and Deletions (Indels), and 144 (8 unique) Copy Number Variants (CNV) across 98 in silico mixed truth sets with a high confidence region covering 2.7 gigabases per mixture. The data is publicly available for use as a benchmarking dataset for somatic short mutation discovery pipelines.

## Introduction

Detecting somatic variation from whole genome sequencing is essential to the understanding and treatment of cancer (1–4). However, discovering somatic short variants (SNVs and Indels) with high precision and sensitivity is still a challenge due to tumor heterogeneity, sequencing artifacts, mapping artifacts and contamination from normal cells (5–7). Many data pipelines and variant calling methods still disagree in a large number of sites, making it unclear which discrepancies are true variants and which are false (8). Even variants that are called by multiple methods are not guaranteed to be true positives. This demonstrates the critical need for high quality benchmarking data that could be used to disambiguate the discrepancies.

Many benchmarking datasets, including ICGC-TCGA DREAM, simulate variation by modifying bases in sequenced reads to known alternate alleles at various allele fractions (9–12). Even with sophisticated modeling, these simulated mutations do not follow the true biological and physical pathways that generate real somatic mutations. As such, synthetic truth data penalize callers that model somatic variation better than the simulations. Other benchmarking datasets often combine germline samples (in silico or in vitro) to simulate heterogeneous tumors. However, germline variation is inherently different than somatic variation in nucleotide substitution frequencies, context, and genomic location frequency (1). In addition, if germline variants are used for benchmarking, somatic variant calling pipelines must disable normal germline filtering. Benchmarking methods that use actual somatic mutations involve expensive validations of individual sites (9). This is limited to a small number of sites and therefore is not powered to make good, unbiased estimates of the performance (5). Another technique of using deeper sequencing (13) as validation, while more sensitive to low allele fraction sites, is no less prone to errors from sequencing, library preparation, or mapping artifacts.

Here we provide a benchmarking dataset of validated somatic mutations in a human colon cancer cell-line with a DNA polymerase epsilon proofreading deficiency (HT115) (14). A known lineage tree structure that was determined using lineage sequencing (LinSeq) (14) is used to validate somatic variation across 11 whole genome HT115 samples, and to construct the high confidence region.

## Using LinSeq to create the benchmark dataset

LinSeq uses imaging technology to record a lineage structure by observing a single cell as it divides for multiple generations. Each of the nodes in the tree (Fig. 1) represents a single cell dividing into two separate sublineages. Once a sufficient number of generations have been observed, several cells are grown up separately and Whole Genome Sequenced to 35X coverage. Variant calling pipelines (see Methods section below) are then run on each of these “leaf” samples and compared with each other to validate the high confidence “branch” variant calls and concurrently define the high confidence regions.

**Fig. 1.**
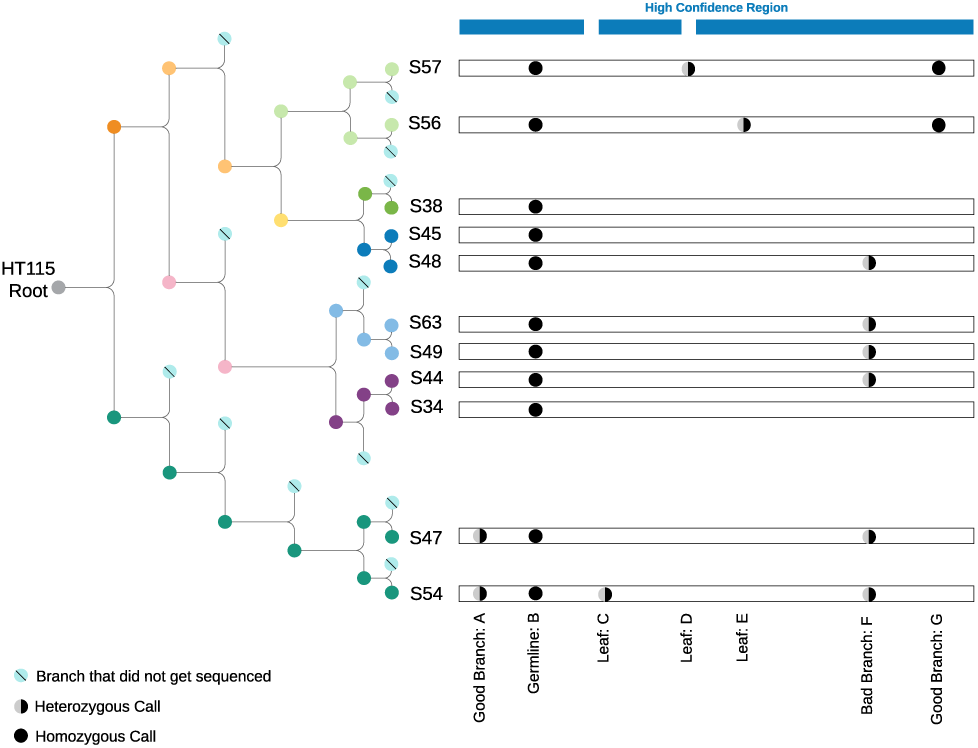
Lineage tree structure for HT115 and an example region of the truth set for S54 as the tumor mixed with S57 as the mixed-in normal. While most of the Good Branch Variants are heterozygous due to single cell bottlenecks, we do observe some Good Branch Variants that are homozygous variant due to loss of heterozygosity events that occurred after the mutation arose or large scale deletions.

Because the different samples contain partially overlapping variants (due to the inheritance structure) the truth set (and high confidence region) depend on the particular tumor-normal pair of samples that are going to be benchmarked against it. Creating the truth set will therefore make use of the inheritance tree and also the calls that were made against all the bulk-sequenced samples.

We also use the LinSeq lineage to create the cases against which the pipeline will be benchmarked. To create a tumor-normal pair at a desired purity, we take three sequenced samples: a “pure tumor” sample and two sister samples that are distant to the tumor. The two sister samples are considered “normal” (relative to the “pure tumor”). We mix (informatically) one of the normal samples with the “pure tumor”to create the case tumor sample, and use the other normal as the case normal, for somatic variant discovery pipelines that are run with the matched normal.

To generate the truth set for each tumor-normal pair of samples, SNVs and Indels are filtered using the lineage structure:

- Germline Variant: a site that is called in all 11 samples. This is either a germline variant of HT115 or a somatic mutation that occurred before the LinSeq process began. For the purposes of the truth set, these sites are considered germline because they occur in both the “tumor” and “normal” sample and therefore the site is not included in the truth set, but remains in the high confidence region. If a somatic variant discovery pipeline calls these variants, they are considered false positives.
- Leaf Variant: a site that is called in only one of the 11 samples. These variants could be real somatic mutations that occurred after the single leaf cell was grown for bulk sequencing, or they could be artifacts that only occurred in one sample by chance. These variants can occur at arbitrary allele fractions because the bulk sequencing process requires growing many cells and can therefore have various subclones within the cell population. Due to these uncertainties, Leaf Variants are removed from the high confidence region. If a variant discovery pipeline makes calls at these sites, they do not count as true positives or false positives.
- Branch Variant: a site that is called in more than one but fewer than 11 samples.
  – Good Branch Variant: a Branch Variant for which all samples with this variant share a single common ancestor, and none of the other samples share that ancestor. A Good Branch Variant that is called in the tumor sample but not the normal is included in the truth set. If a somatic variant discovery pipeline calls these variants, they are considered true positives. These are defined as lineage structure concordant branch variants in the original LinSeq paper (14).
  – Bad Branch Variant: Branch Variants that are not Good Branch Variants. These sites are not included in the truth set, but are not excluded from the high confidence region. If a somatic variant discovery pipeline calls these variants, they are considered false positives.

By only removing Leaf Variants called in the tumor or the mixed-in normal from the high confidence region, we are able to retain sequencing and mapping artifacts in the high confidence region of the truth set (Bad Branch Variants that we are confident are not real somatic mutations).

The true positives generated from this technique are real somatic mutations so pipelines can and should be run exactly as they would be on real samples, with germline and matched normal filtering. These cell-lines also have copy number events that have been validated with the tree structure or are seen in all leaf samples. LinSeq can also be used to validate these CNVs (although at a smaller number of sites than SNVs and Indels, see Table 1). Using these CNV calls we adjust our filtering strategy for SNVs and Indels to account for the expected number of copies in each amplified region.

**Table 1.** Counts of Verified Variants. Counts of events by type. The unique CNVs verified by the lineage tree structure cover 27 megabases with one region at copy ratio 2, one homozygous deletion, and the other six at copy ratio 1.5.

| Type of Variant | Total Sites | Unique Sites |
| --- | --- | --- |
| Verified SNVs | 56974 | 2687 |
| Verified Indels | 6370 | 316 |
| Verified CNVs | 144 | 8 |

## Results

To demonstrate the consistency of this truth set we made all possible pairwise mixtures at three simulated purities^1^, 10%, 20%, and 50%, and ran a somatic variant discovery pipeline on all mixtures. This results in 98 truth sets with various numbers of variants (See Methods section below for how these pairings were determined). Each mixture was analysed with the Mutect2 pipeline in the Genome Analysis ToolKit (GATK) (15) on all 294 case samples at each purity level. For each depth and purity, this results in consistent sensitives and false positives per megabase (Fig. 2). For each subset of purity level and type of variant (SNV or Indel) an overall sensitivity is calculated from the total number of true positives and total positives across all samples.

**Fig. 2.**
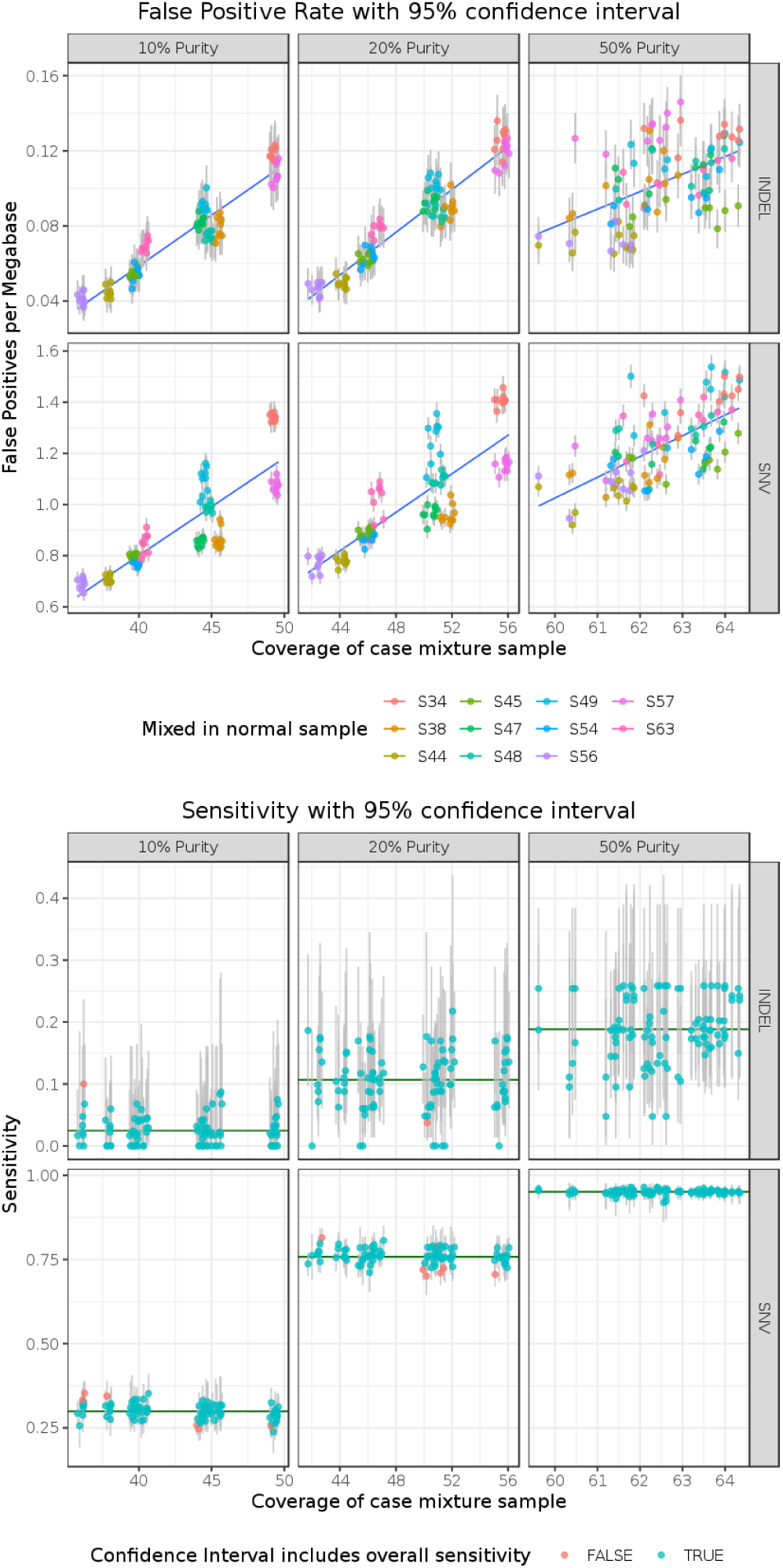
False positive rate and sensitivity from all possible mixtures run with Mutect2. False positive rate clusters by the mixed-in normal sample at lower purities. This is due to variability across samples in quality and false positive rate. Sensitivity follows the expected model for each purity. Green horizontal line denotes overall sensitivity calculated across all mixtures.

A 95% confidence interval for each individual mixture’s sensitivity includes the overall sensitivity derived across all samples for 97% of the mixtures (Fig.2). This demonstrates that the different mixtures measure sensitivities that can reasonably be assumed to come from the same overall distribution. False positive rates (FPR) at 50% purity are consistent given mean coverage of the mixed case sample. (The adjusted R-squared of the linear model predicting FPR from mean coverage of the mixed case sample, type of variant (SNV or Indel), and purity level is .95.) At lower purities, the majority of the reads (80% at 20% purity and 90% at 10% purity) come from the mixed-in normal sample. Quality differences between samples can be seen clearly in the FPR differences given the same overall mean coverage. For example S34 and S57 have different FPR at very similar coverages (Fig.2).

Another way to demonstrate the validity of using the LinSeq lineage tree is to blind ourselves to the tree structure and assess the variants before using the tree as a filter. After applying all filters except the lineage tree (See Methods section below for all filters) we have a list of high quality variants. We can compare the number of these high quality variants that are Good Branch Variants to those that are Bad Branch Variants as another way to check the validity of the lineage tree itself. After removing chromosome 4 due to a large loss of heterozygosity event that occurred in the [S49/S63] branch, and all sites with allele frequency greater than 1% in gnomAD v2.1 (16) to remove common germline variation, a Wilcoxon rank-sum test with continuity correction was performed by ranking each subset of samples based on the number of variants called within that subset (two-sided p-value = 9.0e-8^2^) (Fig. 3). Those subsets that are inconsistent with the tree with the most variants have a subset size of 10 samples and are due to Germline Variants being missed in a single sample (note that the number of Germline Variants is around 5 million). Both Germline Variants called in all 11 samples and Bad Branch Variants, are excluded from the truth set but remain in the high confidence region (since neither artifacts nor true germline variation are true somatic variation).

**Fig. 3.**
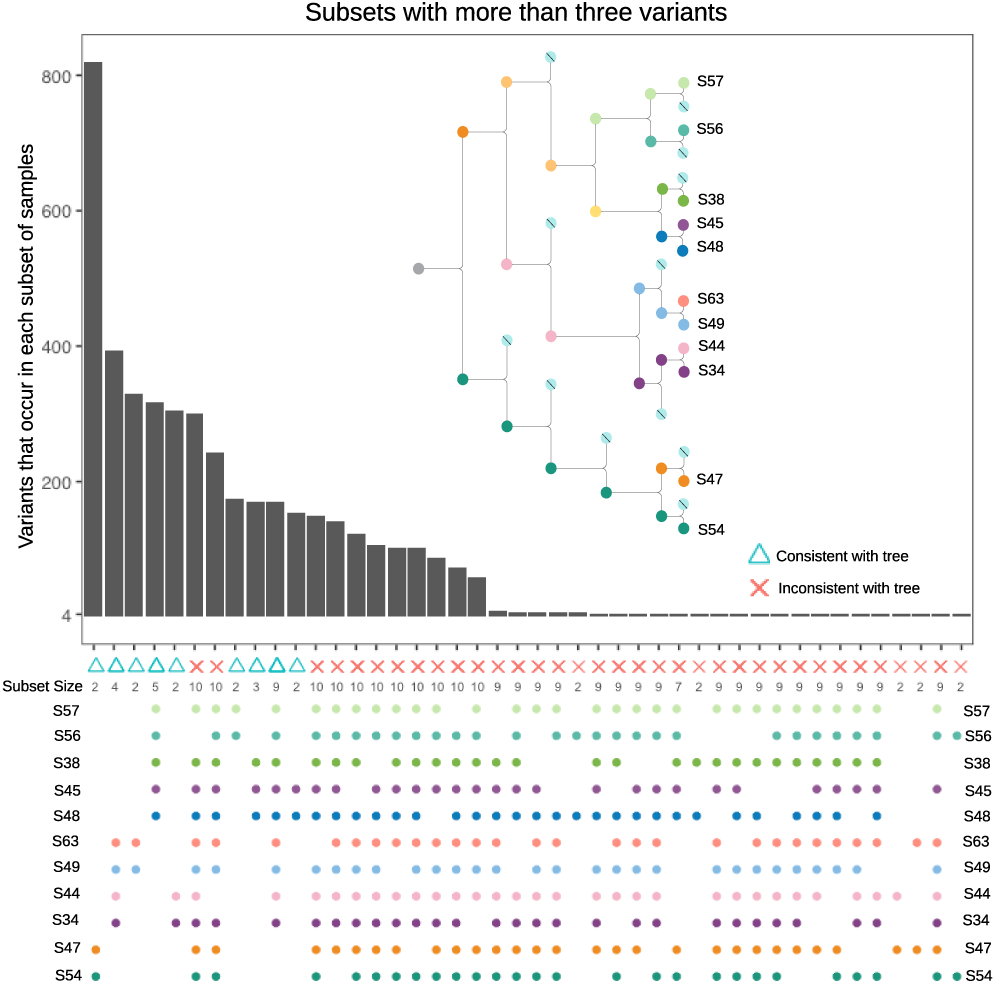
Number of variants called by HaplotypeCaller within each subset. There is a clear separation of subsets that are consistent with the tree structure.

## Methods

### Primary analysis

All data were taken from the original LinSeq experiment by Brody et al. (14). Data can be downloaded from https://www.ncbi.nlm.nih.gov/sra/SRP159787. Samples were reverted and realigned to GRCh38 human reference using the GATK best practice recommendations (15). Reads were aligned with BWA-MEM (17), then duplicate reads were marked using Picard (18). Quality scores were then recalibrated using GATK base quality score recalibrator. Data were stored in per-sample BAM format.

### CNV calling pipeline

GATK’s Model Segments pipeline (19) was used to generate somatic CNV calls. Each sample was run individually with a panel of normals generated from 60 whole genome normal samples from The Cancer Genome Atlas (TCGA) sequenced at the Broad Institute Genomics Platform. The GATK version 4.1.2.0 pipeline was used with the penalty factor set to 5 to combat over-segmentation due to low coverage (samples were sequenced to 35X coverage).

Copy number events were analyzed across all samples to determine whether they follow the tree structure. We obtain copy number amplifications and deletions that follow the tree structure, are called in every sample, or are called in one leaf sample. We find CNVs that don’t follow the tree structure, but it is unclear whether these are due to false positive copy number calls in some samples, or false negatives in other samples.

These copy number calls that did not follow the tree structure are removed from the high confidence region if the tumor sample is not copy neutral in those regions. Copy number calls that only occur in the tumor sample are also blacklisted, since their quality is ambiguous. If the tumor sample is copy neutral, we keep those sites in the high confidence region, since we still see real somatic short variants that follow the tree structure that are unaffected by potential copy number events in other samples (this was confirmed by manual inspection).

We observe 8 unique CNVs that occur within the known lineage tree structure and 646 unique CNVs that occur across all 11 leaf samples. We also see copy neutral loss of heterozygosity in all samples in some regions. This does not affect our short variant analysis since we could still expect somatic mutations to occur in these regions after the loss of heterozygosity, at 50% allele fractions. We do not suspect any genome doubling events in these cells from manual review of the CNV calls.

### SNV and Indel calling pipeline

There are two sections to the SNV and Indel calling pipelines, detecting germline or Good Branch Variants, and detecting Leaf Variants. First, the pipeline for detecting potential true variants is designed to be precise. True variants are expected to follow the diploid assumption (in regions without copy number events) because any mutation went through a single cell bottleneck in the tree structure, and therefore its allele balance should be consistent with being either a hetrozygous or homozygous variant. We only consider variants with good allelle balance as eligible canditates for being true variants. Second, the pipeline for detecting mutations that occurred after the observation of the tree structure (Leaf Variants) is designed to be sensitive even to variants that do not follow the diploid assumption. Each leaf sample was grown up by the lab in order to achieve whole genome bulk sequencing. For that reason there could be subclonal populations in each leaf sample, while the Good Branch variants will be monoclonal. Ambiguity about the source of Leaf Variants is the rationale for excluding these sites from the high confidence region. We use HaplotypeCaller (20) with stringent filters to detect the potential true variants and Strelka2 in somatic mode (21) to detect the potential Leaf Variants.

HaplotypeCaller was run in joint-calling mode, meaning all reads shared local assembly, with all 11 leaf samples as input in order to increase sensitivity. Standard GATK best practices (20) using Variant Quality Score Recalibration (VQSR) was run as an initial filtering strategy. We then filter out any site where any sample has a Genotype Quality (GQ) less than 25. While the majority of sites are expected to have heterozygous genotypes due to single cell bottlenecks, we did not filter out non-heterozygous sites explicitly. Regions with deletions still have homozygous variant genotypes and regions with amplifications were filtered based on the number of copies. Given the copy number call at any site (taken from the CNV calling pipeline described above), any heterozygous site whose Allele Depth (AD) is less than the 1% percentile from *Binomial* 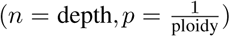 in more than half of the samples is filtered out. This filter is not applied to homozygous variant genotypes, however all other filters are applied. Any site that is “no-called” for any sample is also filtered out. Finally, all of these passing variants that do not follow the expected lineage tree are removed from the list of true variants (but remain in the high confidence region). The known tree structure gives the ability to validate each of the sites across multiple samples separately from individual filters based on read signatures. Genotype information is not taken into account when comparing alleles across the tree structure, so heterozygous and homozygous variant genotypes are all included. The list of passing variants at this point are considered the true somatic variants for each sample (Table 1).

The Strelka2 somatic pipeline is run with each possible pair of leaf samples, one acting as the tumor and the other acting as the matched normal. Thus each of the 11 leaf samples is run through the pipeline as a tumor 10 times, once with each other leaf sample acting as the matched normal, generating up to 10 calls at each site, some of which may be filtered (by Strelka2). If more than half of the calls at any site pass Strelka2’s filters, then the site is included in the potential leaf calls. This is in order to balance the risks of incorrectly calling an artifact a Leaf Variant. If half or more of the calls for one sample at any given site are passing, overall the Strelka2 pipeline believes that call, so we include it as a potential Leaf Variant. The one exception is if all 10 runs are called (filtered or not), the site is included in the potential leaf calls. This is because for many sites called by Strelka2 in all 10 runs even though they looked like sites with borderline evidence in each “tumor-normal” pair, overall they looked like they could be real somatic variation. Finally, when looking at these consolidated filtered calls across all 11 leaf samples, if the call is only found in one sample it is kept as a Leaf Variant and removed from the high confidence region.

### Comparison of SNVs and Indels

In order to assess the quality of the truth set, the Mutect2 (22) pipeline was run on each possible pairing of samples at various mixture rates (to simulate various purities). We then check that sensitivity and FPR for these mixtures run with this variant discovery pipeline are consistent and reasonable. The pipeline was run with a matched normal taken as the sister sample to the mixed-in “normal” sample. This ensures that a similar sample is used as the matched normal to the diluting normal sample, but not the same reads. The Mutect2 pipeline was also run with a panel of normals made from 1000 Genomes Project samples (23) (https://storage.googleapis.com/gatk-best-practices/somatic-hg38/1000g_pon.hg38.vcf.gz), and gnomAD common germline variant filtering (16).

The resulting VCFs were compared within the high confidence region to the truth set using rtgtools VariantEval (24) for both SNVs and Indels to determine Mutect2’s sensitivity and FPR for each mixture.

We take all possible mixtures that will provide true sites at simulated purities of 10%, 20%, and 50%. We have 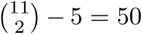 possible pairings since we are picking pairs of samples from the 11 sequenced samples, but we can not use the five pairings of sister samples since there are no variants different between them that we can use the lineage tree to validate ([S57/S56], [S45/S48], [S63/S49], [S44/S34], and [S47/S54]). In addition because S38 has no sister sample, there are no Good Branch mutations that are not also present in the [S45/S48] branch, which removes another pairing (50 1 = 49). Within each pairing there are two possible mixtures (sample 1 as the tumor and sample 2 as the normal or vice versa). This results in 49 * 2 = 98 pairings, each at three purities or 98 * 3 = 294 mixtures.

95% confidence intervals were derived for each mixture’s sensitivity using the exact Clopper-Pearson method, since the number of true positives and false negatives is small enough to use an exact method. 95% confidence intervals for FPR use the asymptotic normal approximation for the binomial distribution, since the larger number of false positives and true negative megabases meets the assumptions for using an asymptotic approximation (Fig. 2).

## Discussion

### Future Work

LinSeq could be repeated on other cancer types and samples to generate other benchmarking datasets, with the potential for testing various wet lab techniques as well. This could be achieved by sequencing each leaf sample twice, once with a control wet lab technique and once with an experimental wet lab technique. Choosing a tumor sample for the root with a matched normal would allow us to disentangle germline variation from somatic mutations that occurred before the observation of the lineage tree. The somatic variant discovery pipelines could also be run with the real matched normal, rather than using a separate leaf sample, removing a complication. Even with the data presented here, more mixtures could be made between three or more samples to simulate tumor heterogeneity.

Deeper sequencing of the samples we have would provide the benefit of getting even lower purity mixtures and more realistic high coverages. It would also give us enough reads to be able to use the same sample as the mixed-in normal and the matched normal, removing another complication in the pipeline and making the purity more realistic. This LinSeq technique could also be useful for generating benchmarking data for CNV callers, although admittedly at much lower number of calls than for short variants. Another similar approach could be to introduce bottlenecks in the cell lineage without using the imaging technology LinSeq uses to determine the lineage tree, but use the variants in the mitochondria to determine the cell lineage (25). This might be a simpler approach in the wet lab, but warrants further investigation if used to generate benchmarking datasets. The hierarchical clustering using distances based on allele fraction from Ludwig et al. (25) was not able to rediscover the known lineage tree on this dataset, perhaps due to the noise introduced from cell colony expansion of each of the 11 samples for whole genome sequencing.

To replicate this experiment as a truly public dataset, not only as sequencing data, but as samples for sequencing, each leaf sample could be continued as cell-lines that could be shared. This would enable further testing of pipelines as well as wet lab techniques. However, the difficulty in this approach would be that somatic mutations would continue to accumulate in these samples over time, meaning that the high confidence region of the truth set would shrink. This might not have a large affect the final benchmarking dataset, but should be considered.

## Conclusion

Using LinSeq our method generates a large number of true somatic variants across the whole genome useful for benchmarking. Since these are actual somatic events there is no need to model features of the variants such as nucleotide context or genomic location. In addition there is no need to change the variant calling pipeline to adapt to deficiencies in the benchmarking dataset such as removing a matched normal, panel of normals, or not filtering known germline variants from the pipeline.

In addition, by using the lineage tree we only have ambiguity in the mutations that occur for a single leaf sample. This means we do not have to remove messy regions that routinely have artifacts, such as homologous regions with mapping errors. Potential false positives that occur in the benchmarking variant calling pipeline are filtered out of the truth set since they do not follow the lineage of the tree, but are still included in the high confidence region of the benchmarking dataset, so they will count as false positives if a variant discovery pipeline calls them.

Finally, by taking CNVs into account, we have better expectations for short variants in regions that have copy number alterations. These regions do not have to be removed from the truth set, but will have varying allele fractions.

We hope that this dataset will be useful for benchmarking new short somatic variant calling pipelines for the general community. All mixed BAMs, truth VCFs, high confidence region interval lists, and benchmarking scripts are available for public use on Terra at https://app.terra.bio/#workspaces/broad-dsp-spec-ops-fc/somatic_truth_data_from_cell_lineage (26). All code to generate these truth datasets are available at https://github.com/meganshand/gatk.

## ACKNOWLEDGEMENTS

We are grateful to Chip Stewart, Takuto Sato, Christopher Kachulis, Mark Fleharty, Madeleine Duran, Sarah Walker, Andrea Haessly, and Sam Friedman for helpful suggestions and feedback on this project.

## AUTHOR CONTRIBUTIONS

Y.E.M. conceived of this study; Y.B. conceived and ran LinSeq and shared the data; J.S. and M.S. analyzed the data and developed the informatic pipelines; M.S. drafted the manuscript; L.L., D.B., Y.F., E.B., and P.C.B. supervised the project. All authors helped to revise the manuscript.

## Footnotes

1 Simulated tumor purity does not mean a single expected allele fraction of all sites in a given mixture. We expect the majority of sites to have an allele fraction that is half of the simulated purity, but due to CNVs and copy neutral loss of heterozygosity events, some sites will be at much lower allele fraction, and others will be at the purity fraction itself.

2 Alternative hypothesis is that the true location shift between the ranks of those subsets that are consistent with the tree and the ranks of those subsets that are inconsistent is not equal to zero. The test statistic is *W* = 1270, sample size for consistent sets is *n* = 9, sample size for inconsistent sets (with at least one variant call) is *n* = 142, estimated difference in location is 301 with a 95% confidence interval of [170, 316].

